# Investigating the in vivo biodistribution of extracellular vesicles isolated from various human cell sources using positron emission tomography

**DOI:** 10.1101/2021.12.29.474459

**Authors:** Zachary T. Rosenkrans, Anna S. Thickens, John A. Kink, Eduardo Aluicio-Sarduy, Jonathan W. Engle, Peiman Hematti, Reinier Hernandez

## Abstract

Noninvasive imaging is a powerful tool for understanding the *in vivo* behavior of drug delivery systems and successfully translating promising platforms into the clinic. Extracellular vesicles (EVs), nano-sized vesicles with a lipid bilayer produced by nearly all cell types, are emerging platforms for drug delivery. To date, the biodistribution of EVs has been insufficiently investigated, particularly using nuclear imaging-based modalities such as positron emission tomography (PET). Herein, we developed positron-emitting radiotracers to investigate the biodistribution of EVs isolated from various human cell sources using PET imaging. Chelator conjugation did not impact EVs size and subsequent radiolabeling was found to be highly efficient and stable with Zr-89 (t_1/2_ = 78.4 h). *In vivo* tracking of EVs isolated from bone marrow-derived mesenchymal stromal cells (BMSCs EVs), primary human macrophages (Mϕ EVs), and a melanoma cell line (A375 EVs) were performed in immunocompetent ICR mice. Imaging studies revealed excellent *in vivo* circulation for all EVs, with a half-life of approximately 12 h. Significantly higher liver uptake was observed for Mϕ EVs, evidencing the tissue tropism of EV and highlighting the importance of carefully choosing EVs cell sources for drug delivery applications. Conversely, the liver, spleen, and lung uptake of the BMSC EVs and A375 EVs was relatively low. We also investigated the impact of immunodeficiency on the biodistribution of BMSC EVs using NSG mice. The spleen uptake drastically increased in NSG mice, which could confound results of therapeutic studies employing this mouse models. Lastly, PET imaging studies in a melanoma tumor model demonstrated efficient tumor uptake of BMSC EVs following intravenous injection. Overall, these imaging studies evidenced the potential of EVs as carriers to treat a variety of diseases, such as cancer or in regenerative medicine applications, and the necessity to understand EVs tropism to optimize their therapeutic deployment.

## Introduction

Extracellular vesicles (EVs) are secreted from nearly all cell types and mediate various physiological and pathological processes, which can mimic the properties of the parental cell from which they are derived.^1,2^ The various types of EVs are generally classified according to their size and biogenesis, including exosomes (~40 - 200 nm), ectosomes (alternatively, microvesicles; ~100 – 1000 nm), and apoptotic bodies (> 500 nm.^3^ The emergence of EVs in biomedical applications has resulted from their ability to efficiently transfer diverse bioactive molecules, such as nucleic acids, proteins, and lipids, to influence the function of target cells.^4^ Additional advantages of EVs in drug delivery applications are their non-immunogenic, inherently biocompatible, biodegradable properties, scalability, and favorable characteristics compared to cell-based therapies.^5^ Due to these properties, EV-based therapies have become an area of intense interest for the treatment of numerous diseases, including cancer and in regenerative medicine applications such as radioprotection.^6–9^

Understanding the *in vivo* biodistribution of EVs is paramount to translating them into successful clinical therapeutics. Tracking exogenous EVs in living subjects has been complicated by their complex composition and small size. Relatively few studies have characterized the *in vivo* fate of EVs using preclinical imaging studies, which have primarily generated confounding results.^10^ The consensus based on previous imaging studies is that EVs are cleared from blood circulation within minutes following injection with uptake confined to the liver, spleen, and lungs.^11^ These studies have led many researchers to question the attractiveness of EVs for clinical applications.

The cell type or tissue sources used to derive EVs can influence their composition and functionality.^12^ Recent studies have found that the activity of multipotent mesenchymal stromal cell (MSC)-based therapeutics is mainly mediated through their release of extracellular vesicles, including EVs containing nucleic acids, proteins, and lipids that accelerate the repair of injured tissues for regenerative medicine applications.^13–15^ MSCs, under investigation in numerous clinical trials, often isolated from human tissues such as bone marrow (BMSC), adipose, or umbilical cords, can all be used to isolate EVs for regenerative medicine applications.^16,17^ Macrophage-derived EVs (Mϕ EVs) have also been investigated as regenerative medicine therapeutics or cancer therapies.^18–21^ Despite the various and vast applications of EVs, the *in vivo* biodistribution remains relatively unexplored, particularly using nuclear medicine imaging modalities such as positron emission tomography (PET).

Herein, we investigated the *in vivo* fate of EVs isolated from different parent cell types using PET imaging, a highly sensitive, quantitative, noninvasive technique offering limitless tissue penetration and easy scalability from preclinical to large animal or human investigations. EVs isolated from three distinctive cell types, MSC, macrophages, and cancer cells, were surface-conjugated with a chelator (deferoxamine, Df) and radiolabeled efficiently and stably with Zr-89 (t_1/2_ = 74.8 h). Subsequently, PET imaging studies compared the biodistribution of the different EVs in immunocompetent and immunodeficient mice. Through describing the *in vivo* fate of EVs from diverse cell sources, our in vivo tracking studies bring attention to the importance of the organotropic properties of EVs and their potential impact on their development as therapeutics for various diseases.

## Materials and methods

### Isolation and Cultivation of cells

Bone marrow-derived MSCs (BMSCs) were first isolated as previously described from the bone marrow (BM) of healthy human donors.^22^ Briefly, leftover BM cells from transplant donors were washed with PBS, and mononuclear cells were isolated using Ficoll-Paque Plus density gradient separation. ACK lysis buffer was used to lyse red blood cells with a 3 min incubation as needed. Mononuclear cells were suspended in α-minimum essential medium supplemented with 10% fetal bovine serum, 1% penicillin-streptomycin, 1x nonessential amino acids, and 4 mM L-glutamine. The identity of MSCs was confirmed using flow cytometry [CD90 (+), CD105 (+), CD34 (-), CD45 (-)]. Passage 4 to 8 MSCs were used for EVs isolation (BMSC EVs).

Monocytes were isolated from human peripheral blood mononuclear cells (PBMCs) obtained from the blood of healthy stem cell donors as described previously.^7^ Briefly, PBMCs were separated using density gradient separation with Ficoll-Paque Plus, and RBCs were lysed using ACK lysis buffer. Platelet contamination in cell suspension was reduced by centrifuging the cell suspension at 300-700 rpm for 10 mins. CD14^+^ monocytes were collected in the resuspended cell suspension by incubating with anti-human CD14 microbeads (Miltenyi Biotec) for 15 mins at 4°C and separated using a MACS Pro Separator (Miltenyi Biotec). Monocytes were cultured using Iscove’s Modified Dulbecco’s media supplemented with 10% human serum blood type AB, 1x nonessential amino acids, 4 mM L-glutamine, 1 mM sodium pyruvate, and 4 μg/mL recombinant human insulin. For differentiation into macrophages, monocytes were cultured for 7 days at 37°C with 5% CO_2_ without cytokines.

Human malignant A375 melanoma cells (ATCC; Manassas, VA) were cultured in Dulbecco’s Modified Eagle’s Medium (DMEM) supplemented with 10% FBS per vendor recommendations.

### Ultracentrifugation of EVs from cells

EVs were isolated from the various cells (BMSCs, macrophages [Mϕ], A375) types were grown in complete media as described above using sequential ultracentrifugation. Cells were washed twice with PBS and media replaced with MSC serum-free media (SFM; StemPro A103332-01, ThermoFisher Scientific). Cells were incubated for 18-24 h in SFM, and the collected conditioned media was centrifuged at a low-speed spin (2000 x g at 4°C for 20 minutes) to remove any detached cells and cell debris. The supernatant was then ultracentrifuged for 2 hours using Optima™ L-80XP Ultracentrifuge (Beckman Coulter) at 100,000 x g at 4°C. Purified EVs (BMSC EVs, Mϕ EVs, and A375 EVs) were then resuspended in PBS and stored at −80°C until further use.

### EVs conjugation

EVs were conjugated to p-SCN-Bn-Deferoxamine (Df; Macrocyclics) for PET imaging. In a typical reaction, EVs were conjugated to Df using isothiocyanate chemistry by dissolving Df in anhydrous DMSO and added to EVs in PBS that was pH adjusted to 8.5-9. The reaction was conducted at room temperature for 1-2 h. The conjugated EVs were purified using a PD-10 size exclusion chromatography column.

### EVs radiolabeling

Conjugated EVs (Df-BMSCs, Df-Mϕ EVs, or Df-A375 EVs) were radiolabeled with Zr-89 for PET studies (^89^Zr-Df-BMSCs, ^89^Zr-Df-Mϕ EVs, or ^89^Zr-Df-A375 EVs). For radiolabeling, 1-2 mCi (37-74 MBq) of Zr-89 was added to 1 M HEPES buffer, and Df conjugated EVs were added. The reaction mixture was incubated at 37°C for 1 h and labeled EVs purified using a PD-10 size exclusion chromatography column (GE Healthcare). Instant thin layer chromatography (iTLC) was used to determine the radiolabeling efficiency and radiostability in PBS and 50% serum.

### Animal studies

All animal studies were conducted on a protocol approved by the Institutional Animal Care and Use Committee at the University of Wisconsin-Madison. Hsd:ICR (CD-1®) [ICR] mice were purchased from Envigo. C57BL/6 and NSG mice were purchased from Jackson Laboratory and bred at the University of Wisconsin-Madison.

### PET/CT imaging studies

PET/CT images were acquired using an Inveon μPET/CT scanner. CT images were captured using the following parameters (80 kV, 900 uA, resolution of 105 μm). Mice were intravenously injected with approximately 5.55 MBq – 7.4 MBq (150 μCi – 200 μCi). Static PET images were acquired at various timepoints post injection (2-3 h, 24 h, 48 h, 72 h, and/or 144 h) and reconstructed using an OSEM3D/MAP algorithm. The Inveon Research Workstation software was used to perform quantitative region of interest analysis of the PET images, with values reported in percent injected activity per gram of tissue (%IA/g). After the final imaging time point, the mice were euthanized, and the major organs were collected for *ex vivo* biodistribution studies. A Wizard 2 (Perkin Elmer) gamma counter was used to quantify tracer uptake in organs. Biodistribution data was calculated in percent injected activity per gram of tissue (%IA/g) for all organs of interest.

### Murine B78 melanoma model

To establish B78 melanoma tumor grafts, C57BL/6 mice were intradermally injected with 2×10^6^ B78 cells on the lower right flank of the mice. Mice were used for PET/CT imaging studies once tumors reached 200-300 mm^3^, measured by tumor volume = (1/2) x length x width^2^.

## Results and discussion

### EVs Isolation, radiolabeling, and PET imaging

Understanding the *in vivo* biodistribution and pharmacokinetics, which the parental cell type may influence, is fundamental to successfully developing EV-based therapeutic agents (**Figure 1a**). In addition, properly understanding these properties is key to establishing EVs dosing route and regimen, understanding their mechanism of action, and exposing potential off-target effects leading to toxicity. To this end, we chose to investigate the *in vivo* properties of EVs derived from three distinctive cell sources, human BMSC, Mϕ, and A375 melanoma cells. As shown in **Figure 1b**, the average size of the isolated EVs was comparable (BMSC EVs: 136 ± 2.6 nm, Mϕ EVs: 117.6 ± 0.4 nm, and A375 EVs: 123.8 ± 3.6 nm) following sequential ultracentrifugation. Thus, differences in the *in vivo* biodistribution would be attributed mainly to the cell type used for EVs isolation.

**Figure 1.**
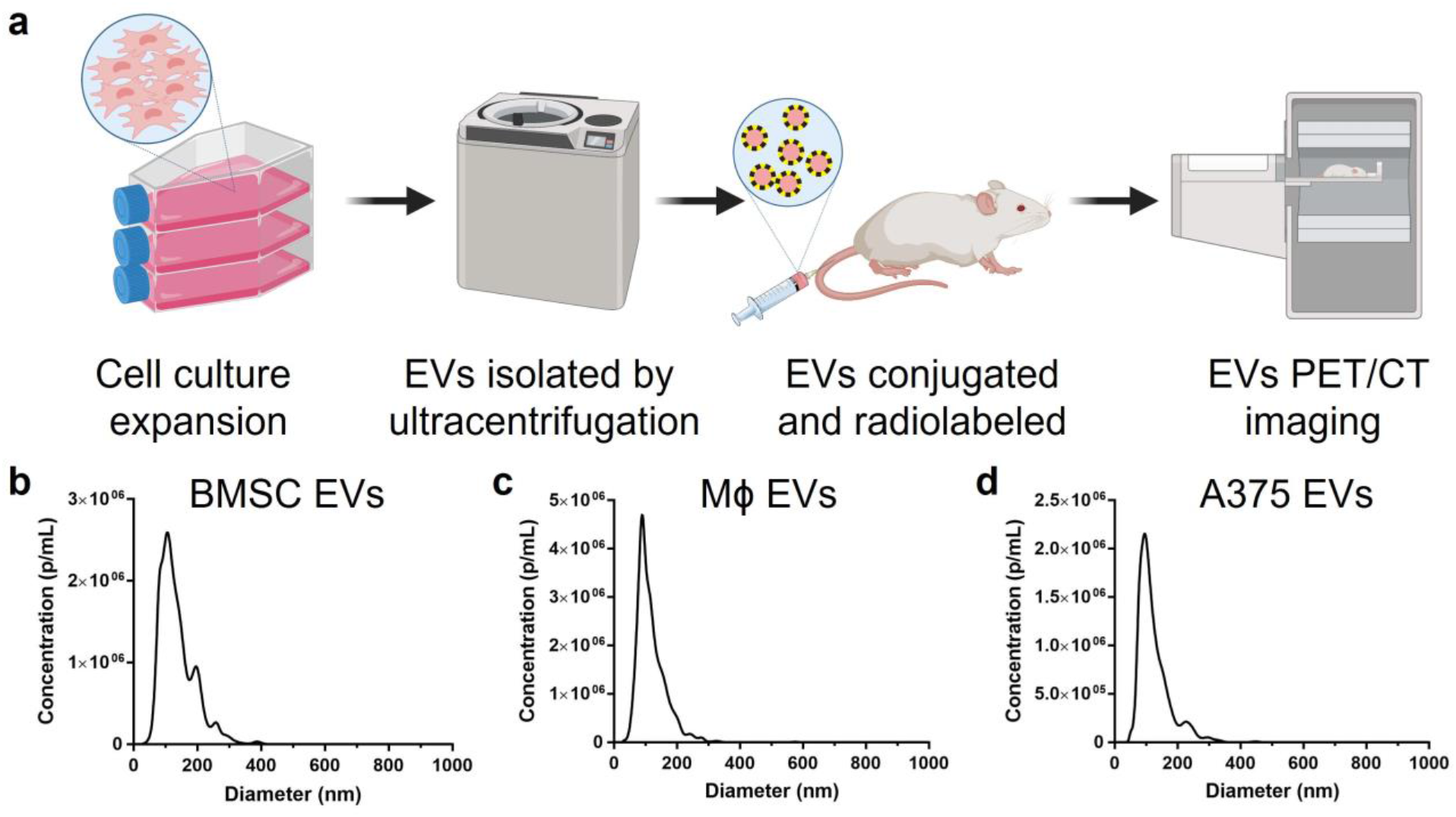
The *in vivo* biodistribution of EVs isolated from various human cell sources was investigated using positron emission tomography (PET). (**a**) Cells were isolated and cultured for EVs isolation using sequential ultracentrifugation. EVs were then conjugated with a deferoxamine chelator and radiolabeled with Zr-89 (t_1/2_ = 78.4 h) for PET studies. Nanoparticle tracking analysis found that EVs isolated from (**b**) BMSC EVs, (**c**) Mϕ EVs, and (**d**) A375 EVs were all similarly sized, with an average diameter of approximately 110-140 nm.

Ideal labeling methods for imaging studies must not disrupt the physicochemical properties of the traced agent while providing high sensitivity and *in vivo* stability. Additionally, the labeling method will dictate the imaging modality and thereby the interpretation of EVs localization *in vivo*.^23^ We chose to stably conjugate and chelate EVs with a positron-emitting isotope for PET/CT imaging to investigate the biodistribution of EVs from different cell types, which may be critical when using EVs as drug delivery systems for tissue or organ targets. Using PET as our imaging modality allows minimal EVs modification because of its high detection sensitivity, enabling longitudinal tissue uptake quantification. For imaging studies, deferoxamine (Df; Df-BMSC EVs, Df-Mϕ EVs, or Df-A375 EVs) was conjugated to EVs using isothiocyanate chemistry and subsequently radiolabeled with Zr-89 (^89^Zr-Df-BMSC EVs, ^89^Zr-Df-Mϕ EVs, or ^89^Zr-Df-A375 EVs; **Figure 2a**). The influence of chelator conjugation on EVs size was analyzed using nanoparticle tracking analysis, which found comparable mean (116 nm vs. 118 nm) and mode (80 nm vs. 81 nm) diameters for both BMSC and Df-BMSCs EVs (**Figure 2b**). EVs were successfully radiolabeled with Zr-89 with reported radiolabeling efficiency for 1 h reactions of 88.3 ± 1.4% and 89.6 ± 0.3%, which corresponded to specific activity per particle of 4.19 × 10^−9^ MBq/EVs and 4.13 × 10^−9^ MBq/EVs for ^89^Zr-Df-BMSCs EVs and ^89^Zr-Df-Mϕ EVs, respectively (**Figure 2c-d**). Importantly, we determined the stability of the radiolabeled EVs in PBS and 50% mouse serum (**Figure 2e**). Using ^89^Zr-Df-BMSCs EVs as a representative sample, labeled EVs showed long-term stabilities of 91.6 ± 5.0% and 85.0 ± 2.9% after 72 h of incubation in PBS and 50% mouse serum, respectively. Thus, we demonstrated that EVs can be conjugated without altering physical properties and that they could be efficiently and stably radiolabeled with Zr-89 for *in vivo* PET studies.

**Figure 2.**
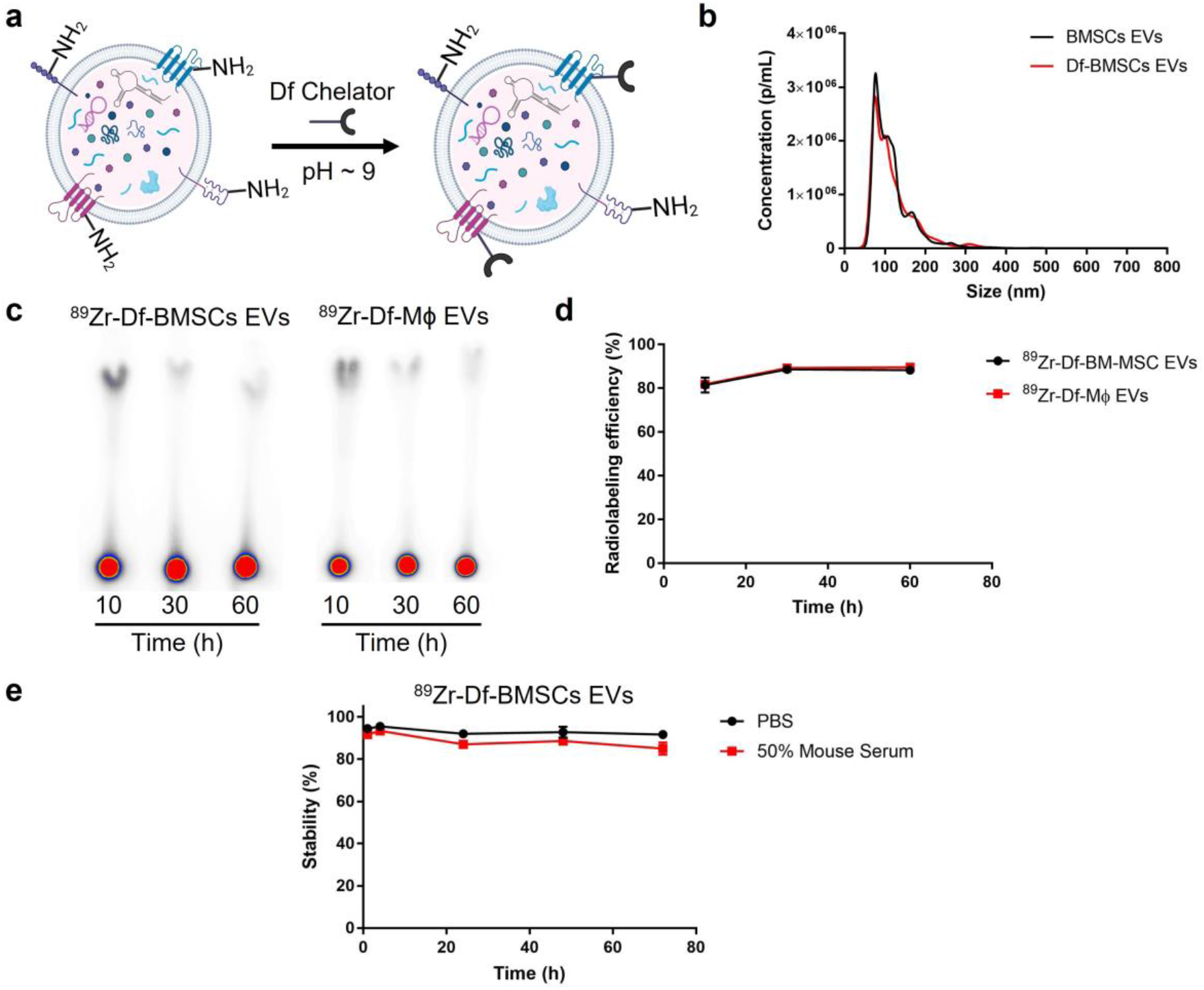
Conjugation and radiolabeling of EVs for PET. (**a**) Schematic showing the conjugation of the chelator (Df) to the surface of EVs. (**b**) Nanoparticle tracking analysis showed no differences in the size distribution of BMSC EVs and Df-BMSC EVs after conjugation. (**c**) Representative instant thin layer chromatography used to determine the (**d**) radiolabeling efficiency, which exceeded 85% (n=3). (**e**) Stability studies of ^89^Zr-Df-BMSC EVs showing that radiolabeling was stable in PBS and 50% mouse serum up to 72 h following incubation (n=3).

Given the comparable properties of native and conjugated EVs and their efficient and stable radiolabeling with Zr-89, we investigated the biodistribution of EVs from different cell sources following intravenous injection using PET/CT imaging in healthy ICR mice (**Figure 3**). Volume of interest (VOI) analysis of serial PET/CT images quantified uptake in major organs of interest (**Figure 4a-g**). Prolonged circulation half-lives for all EVs were observed, and they were all largely cleared from the blood at 72 h post-injection (p.i.). Fitting the time-activity curves of the heart showed blood circulation half-lives of 12.4 ± 0.3 h, 14.4 ± 1.1 h, and 12.3 ± 0.6 h for ^89^Zr-Df-BMSC EVs, ^89^Zr-Df-Mϕ EVs, and ^89^Zr-Df-A375 EVs, respectively. Interestingly, these results contrast previous studies reporting EVs blood circulation of less than 5 min.^24^ The primary clearance route for all the EVs was hepatic, with liver uptake of ^89^Zr-Df-BMSC EVs, ^89^Zr-Df-Mϕ EVs, and ^89^Zr-Df-A375 EVs reaching maximums at 2 h p.i. of 6.9 ± 0.9 %IA/g, 26.3 ± 4.4 %IA/g, of 7.7 ± 0.3 %IA/g, respectively. The liver uptake for all EVs then slowly decreased throughout the imaging study. The most notable difference in distribution between the imaged EVs was seen in liver uptake of ^89^Zr-Df-Mϕ EVs, which ranged from 3.3-4.5-fold higher across imaging timepoints than either ^89^Zr-Df-BMSC EVs or ^89^Zr-Df-A375 EVs. Interestingly, no other notable differences between the EVs uptake in the spleen, lung, kidney, muscle, or bone were observed. All EVs had minimal uptake in the lungs or kidneys, suggesting no apparent aggregation or destabilization following EVs administration. Uptake in blood-rich organs, including the lung and kidney, decreased throughout the imaging study in conjunction with the blood clearance. Notably, minimal radioactivity, which plateaued immediately following injection, was detected in the bones indicating the low levels of bone-seeking “free” Zr-89, confirming the *in vivo* stability of the radiolabeled EVs. *Ex vivo* biodistribution studies following the final imaging time point at 72 h p.i. were performed to validate PET/CT results and to further quantify EVs uptake in other major organs of interest (**Figure 4h**). In line with the imaging data, for all EVs, the most prominent uptakes were found in the liver (^89^Zr-Df-BMSC EVs: 6.45 ± 0.55 %IA/g, ^89^Zr-Df-Mϕ EVs: 29.04 ± 3.41 %IA/g, ^89^Zr-Df-A375 EVs: 8.92 ± 0.52 %IA/g) and spleen (^89^Zr-Df-BMSC EVs: 5.93 ± 0.60 %IA/g, ^89^Zr-Df-Mϕ EVs: 10.62 ± 4.26 %IA/g, ^89^Zr-Df-A375 EVs: 6.76 ± 0.52 %IA/g). The uptake of the imaged EVs in the other major organs correlated to the VOI analysis and appeared non-specific, with all values less than 3.5 %IA/g. Based on these results, EVs exhibited excellent *in vivo* circulation and generally had similar biodistribution. Nonetheless, significant differences in the EVs biodistribution were observed for some specific organs, highlighting the importance of considering the cell source used for EVs production when developing therapeutic agents.

**Figure 3.**
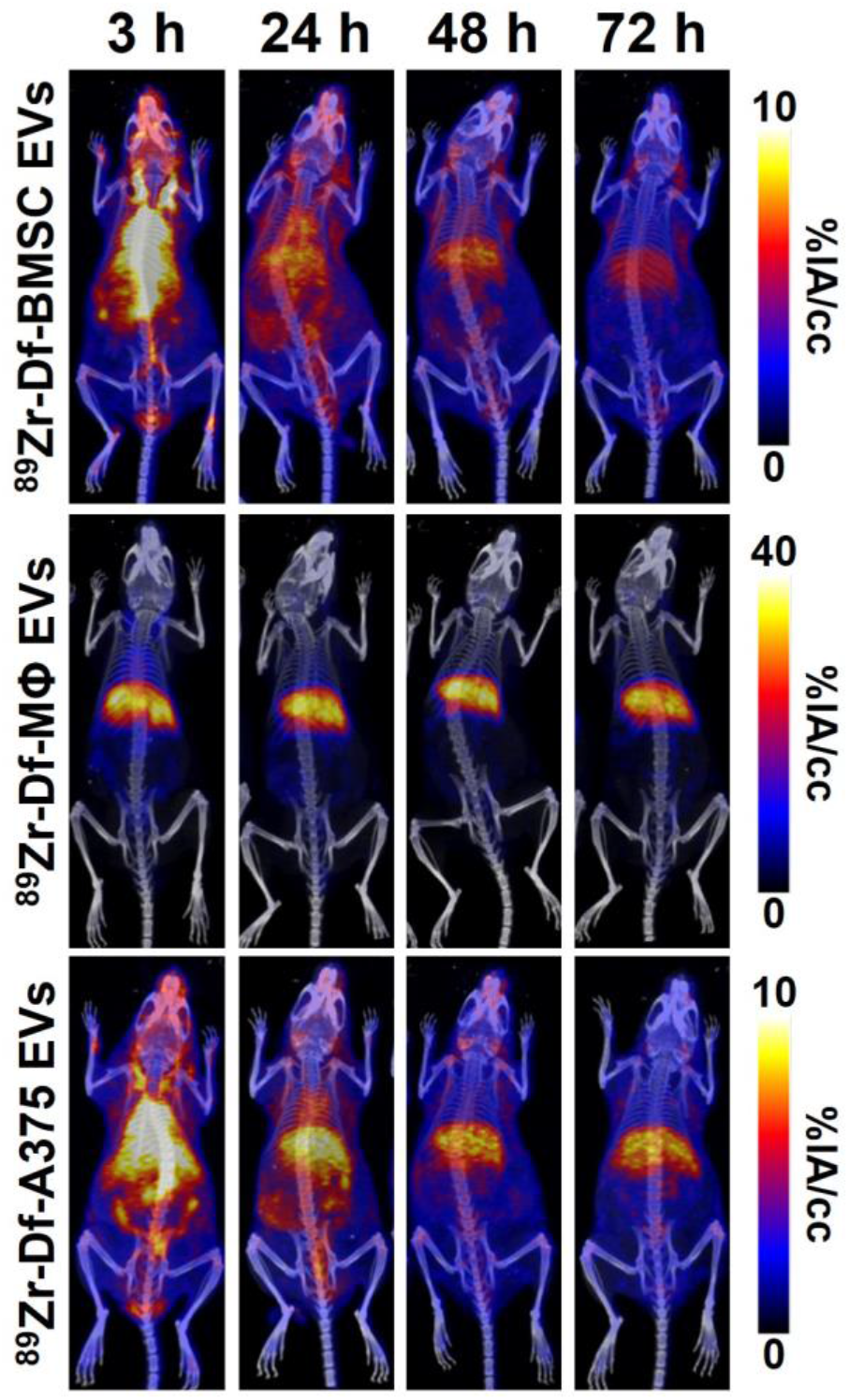
PET imaging of EVs in ICR mice. Following conjugation and radiolabeling, ^89^Zr-Df-BMSC EVs, ^89^Zr-Df-Mϕ EVs, and ^89^Zr-Df-A375 EVs were intravenously injected in ICR mice and serial PET/CT images were acquired at various times post-injection. Maximum intensity projections of the fused PET/CT images are shown.

**Figure 4.**
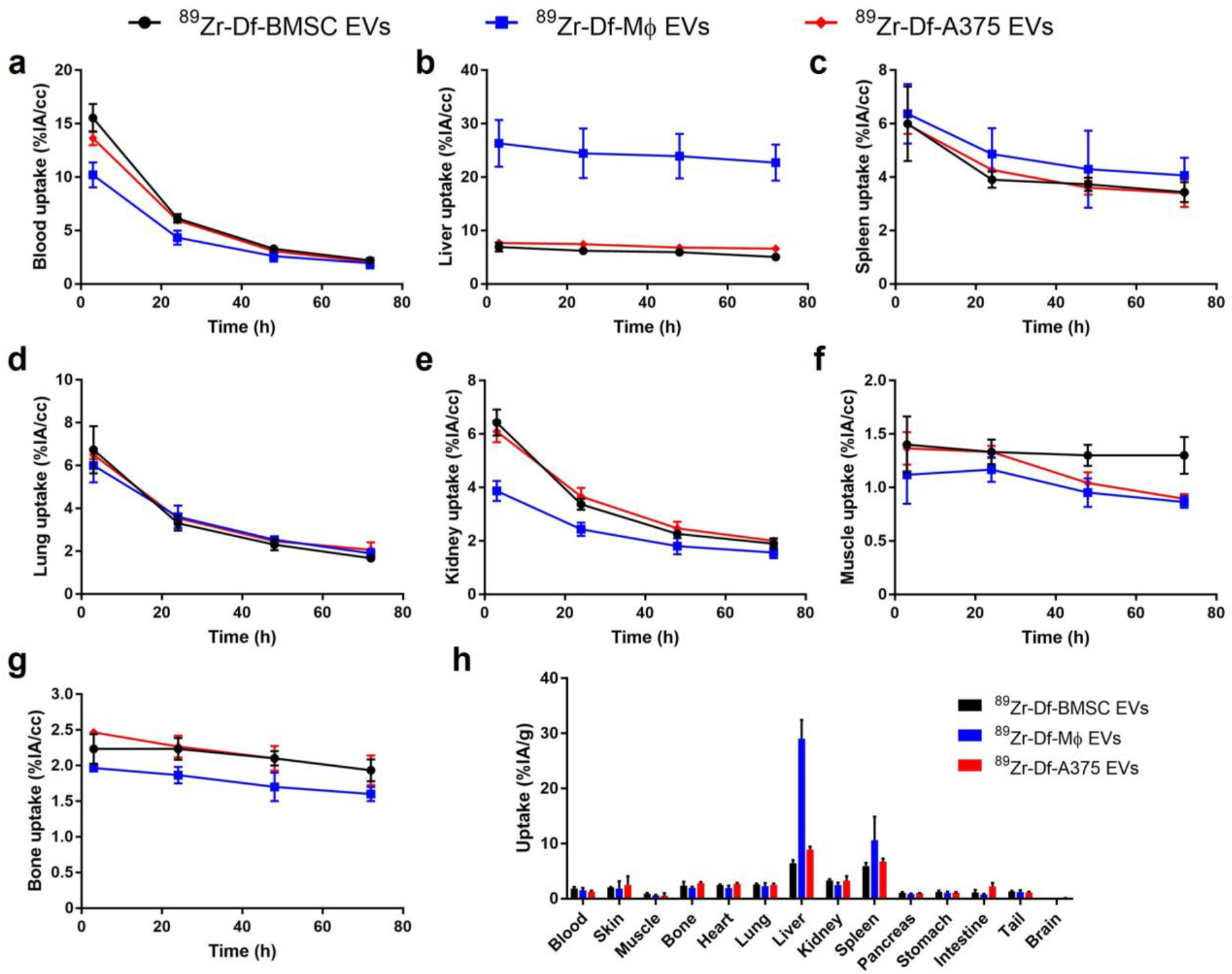
Quantitative PET analysis of EVs distribution in healthy mice. Volume of interest quantification of PET images was performed on (**a**) blood, (**b**) liver, (**c**) spleen, (**d**) lung, (**e**) kidney, (**f**) muscle, and (**g**) bone at each imaging timepoint (n=3). (**h**) *Ex vivo* biodistribution studies quantified uptake in the major organs following the final imaging timepoint at 72 h post-injection (n=3).

In this study, EVs were labeled by covalent modification using the primary amine groups present on surface proteins, which affix the imaging label to the EVs surface to limit background signal and can be utilized for fluorescent (FL) or nuclear imaging applications.^25–28^ Other strategies have been investigated for labeling EVs, with the most common strategy being membrane integration. Membrane integration labeling most frequently employs lipophilic molecules, such as dialkylcarbocyanine dyes (DiD, DiR, etc.), PKH dyes (PKH26, PKH67, etc.), or radioactive labels that integrate into the EVs lipid bilayer non-specifically.^29–35^ Using lipophilic membrane incorporating dye can result in the exchange between EVs and cells in the subject that confounds image interpretation. In addition, although simple and easy, this labeling method is limited by background from dissociated labels, induction of EVs clumping, and inability to distinguish between lipid proteins and micelles. Using this labeling strategy, previous studies investigated the influence of cell source, route of administration, and targeting on EVs biodistribution.^36^ However, the FL imaging employed in their study is neither quantitative nor able to reliably track EVs biodistribution *in vivo* with accurate spatiotemporal resolution because of the limited penetration of light in tissues, poor label stability, and low sensitivity requiring the loading of a large number of dyes onto the EVs. Thus, FL imaging only showed minor differences in EVs biodistribution isolated from different cell types. The advantages and limitations of various imaging labeling strategies must be carefully considered in any application with conditions optimized to attain meaningful results for understanding the *in vivo* biodistribution of EVs.

In general, we observed similar trends in tissue biodistribution between the various EVs; however, the cell type source employed could induce drastic changes in distribution to organs like the liver and spleen. ^89^Zr-Df-BMSC EVs and ^89^Zr-Df-A375 EVs showed similar biodistribution with minimal uptake in the liver, kidneys, spleen, and lungs. On the other hand, ^89^Zr-Df-Mϕ EVs showed fast and prominent accumulation in the liver, with minimal decline in uptake over the 72 h observation period. The contrasting results in the liver and spleen could be explained by an inherent tropism of the Mϕ-derived EVs to these macrophage-rich tissues. As such, EVs organotropism is likely influenced by the composition of their membranes, favoring organs that are hosts to the parental cells from which they are derived. Studies have described similar cell tropisms in biological processes or drug delivery applications using cell carriers, a property that may be likely passed to the cellular-derived EVs.^37–39^ Thus, this intrinsic tropism of cellular sources of EVs is an essential factor to consider when optimizing the design of drug delivery platforms for a specific disease.

### Effects of immune status on EVs Biodistribution

Research applications often require using immunodeficient mice to establish human cancer models or investigate the efficacy of cell therapies.^7^ Therefore, we used serial PET/CT imaging to examine the impact of a dysfunctional adaptive immune system on the biodistribution of ^89^Zr-Df-BMSC EVs using healthy immunocompromised NSG mice lacking T-cells, B-cells, and NK cells (**Figure 5a**).^40^ As we previously observed, ^89^Zr-Df-BMSC EVs had prolonged circulation and minimal uptake in the lungs or liver in NSG mice. Using VOI analysis (**Figure 5b**), we found that ^89^Zr-Df-BMSC EVs were predominately cleared from the blood by 72 h p.i., with a blood circulation half-life of 11.6 ± 0.9 h. High spleen uptake of ^89^Zr-Df-BMSC EVs in the NSG mice was evident from the initial imaging timepoint. Previous studies have found similarly high spleen uptake of human monoclonal antibodies in NSG mice, attributed to Fc region – Fc receptor binding on myeloid cells in these mice with low to no endogenous immunoglobulins.^41^ Moreover, Fc region – Fc receptor interaction and overall spleen uptake were correlated to the degree of glycosylation on the Fc region of the antibody.^42^ Glycans are a fundamental component on the surface of exosomes, and the importance of glycosylation status on the observed spleen uptake of NSG mice should be investigated in future studies.^43,44^ The ^89^Zr-Df-BMSC EVs exhibited hepatic excretion with liver uptake plateauing at 24 h at 6.20 ± 1.01 %IA/g. As in immunocompetent mice, minimal uptake of free Zr-89, peaking at 2.0 ± 0.5 %IA/g 24 h p.i., was observed in the bones at all time points. *Ex vivo* biodistribution studies after the final imaging time point at 144 h p.i. determined exosome uptake in the major tissues of interest (**Figure 5c**). The spleen had the most prominent uptake of ^89^Zr-Df-BMSC EVs (55.6 ± 11.2 %IA/g), which could be attributed to the EVs glycosylation status.^44^ The significant difference between the spleen uptake quantified from VOI analysis and *ex vivo* biodistribution studies can be explained by partial volume effects associated with the small spleen size in NSG mice and limited spatial resolution of PET.^45^ The uptake of ^89^Zr-Df-BMSC EVs in the other major organs, however, correlated to the VOI analysis and appeared non-specific, with only the liver having uptake greater than 5 %IA/g. This imaging study confirmed the excellent *in vivo* circulation of MSC-derived EVs, but the observed prominent spleen uptake in NSG mice reaffirmed the necessity for performing distribution studies in specific mouse strains. These results are critical to inform therapeutic studies involving MSC-derived EVs that require utilization of NSG mice, as divergent biodistribution patterns in this mouse strain might confound interpretation of experimental results.

**Figure 5.**
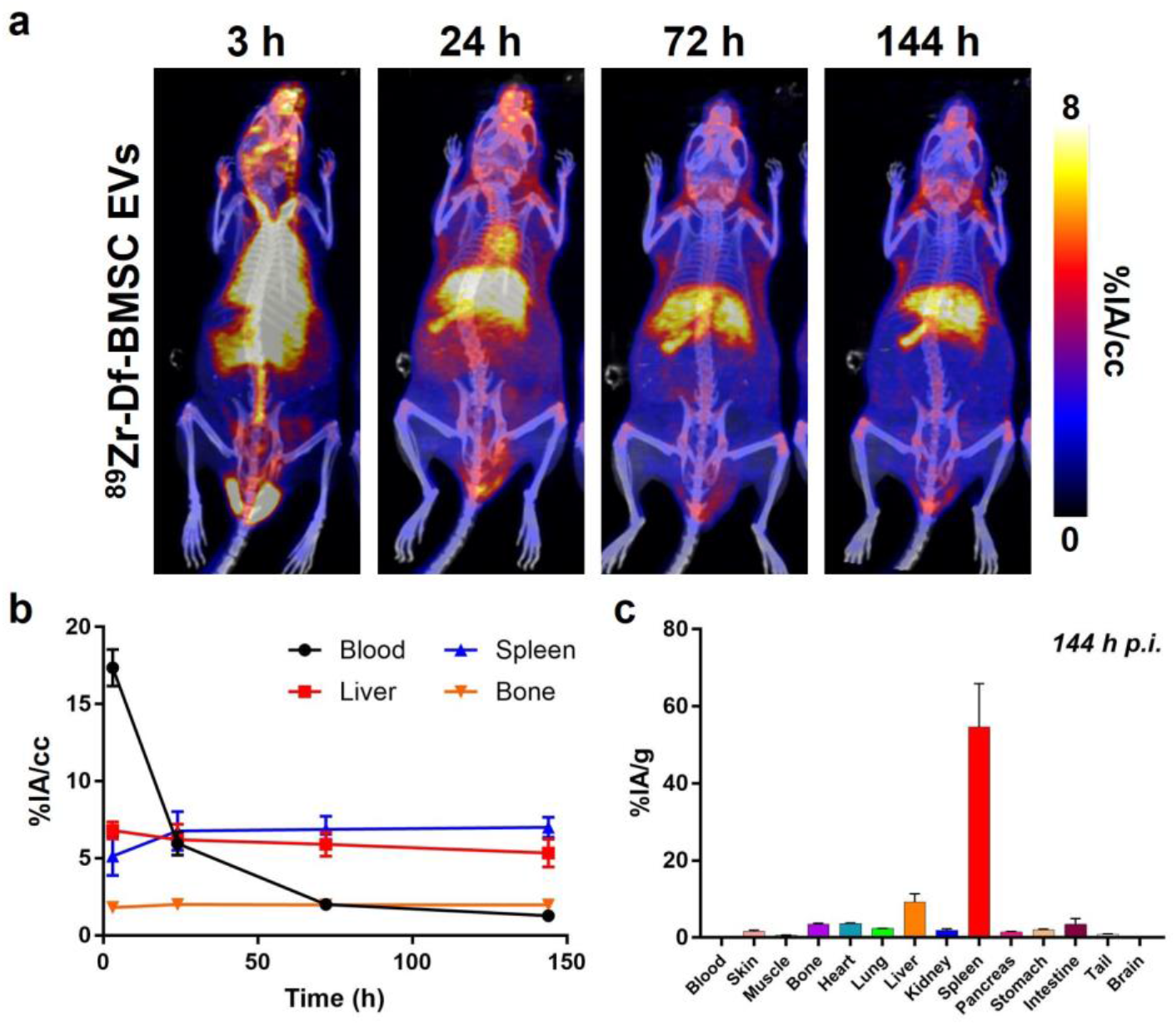
PET imaging of EVs in immunocompromised NSG mice. (**a**) Serial maximum intensity projection PET/CT images of healthy NSG mice intravenously injected ^89^Zr-Df-BMSC EVs. (**b**) Volume of interest quantification of ^89^Zr-Df-BMSC EVs at various timepoints post-injection (n=3). (**d**) *Ex vivo* biodistribution studies of ^89^Zr-Df-BMSC EVs following the final imaging timepoint at 144 h post-injection (n=3). Marked liver and spleen uptake were observed in these mice. Incongruences in spleen uptake between imaging and *ex vivo* results are likely attributed to partial volume effects.

### PET imaging of EVs in cancer model

Due to the potential of MSC-derived EVs and other EVs as cancer therapies,^46^ we also explored the tumor uptake and distribution of BMSC EVs in a syngeneic B78 melanoma mouse tumor model. PET/CT images were acquired after intravenously injecting ^89^Zr-Df-BMSC EVs in mice bearing B78 tumors and quantified tissue uptake with VOI analysis (**Figure 6**). The circulation of ^89^Zr-Df-BMSC EVs was prolonged up to 48 h p.i., with a circulation half-life of 12.0 ± 1.7 h. Notably, the tumor could be visualized at all time points, although contrast decreased at the final time point at 144 h p.i. ^89^Zr-Df-BMSC EVs tumor uptake peaked at 24 h p.i. with a value of 5.3 ± 0.5 %IA/g. The tumor uptake of ^89^Zr-Df-BMSC EVs can be attributed to the enhanced permeability and retention effect.^47^ Relatively low uptake in the liver and spleen was found, which reached maximums at 3 h p.i. when blood-pool activity was highest and then was slowly eliminated up to 144 h p.i. of the radiotracer. *Ex vivo* biodistribution studies after the terminal imaging time point at 144 h p.i. quantified the uptake of ^89^Zr-Df-BMSC EVs in the major organs of interest (**Figure 6c**). The highest uptake of ^89^Zr-Df-BMSC EVs was observed in the spleen (12.8 ± 2.6 %IA/g) and liver (11.2 ± 2.2 %IA/g). In comparison, the B78 tumor uptake of 89Zr-Df-BMSC EVs was 3.6 ± 1.8 %IA/g. This imaging study demonstrates the efficient delivery of systemically administered EVs to tumors, corroborating the potential of MSC-derived EVs as carriers of drugs or engineered bioactive molecules for cancer therapy. Engineering EVs for cancer therapies are among the most promising therapeutics undergoing clinical trials (ClinicalTrials.gov identifiers: NCT05156229, NCT04592484).^48–50^ Utilizing PET approaches in humans would be ideal for understanding the biodistribution of EVs over time, which is an essential requirement for successfully developing drug delivery platforms. Understanding the pharmacokinetic profile through noninvasive imaging informs tumor uptake and off-target targeting toxicity concerns, which can be used to design more effective, safer therapeutics and accelerate translation. The tumor accretion could be further improved by engineering EVs to express molecular binders targeting tumor antigens, which may further increase their specificity and potency as drug delivery platforms. This approach will require an in-depth understanding of the *in vivo* distribution profile of the engineered EVs using noninvasive PET methodologies like the ones described herein.

**Figure 6.**
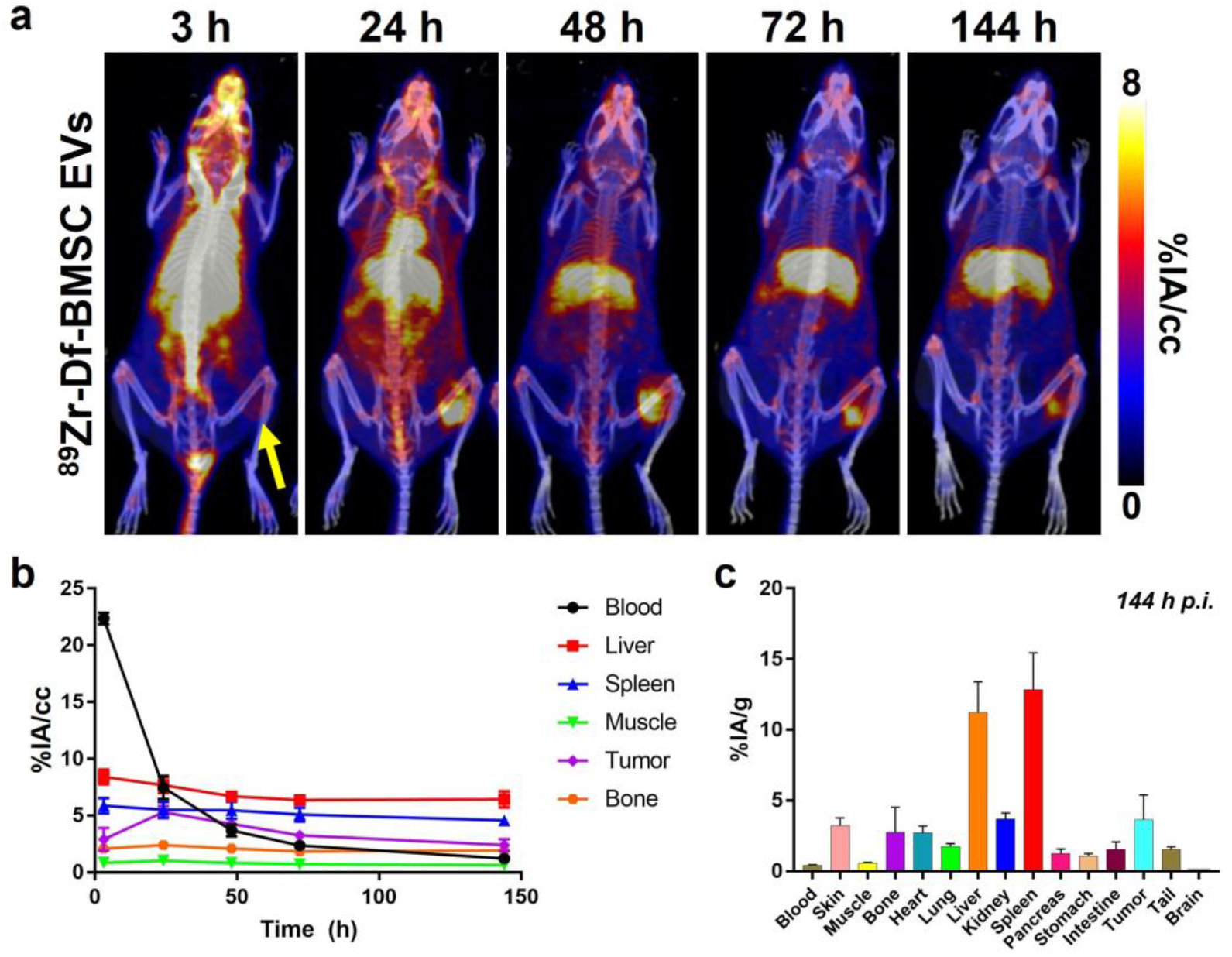
PET imaging of ^89^Zr-Df-BMSC EVs in a syngeneic B78 melanoma mouse model. (**a**) Serial maximum intensity projection PET images in B78 tumor bearing mice intravenously injected ^89^Zr-Df-BMSC EVs show marked tumor uptake that peaked at 24 h post-injection (p.i.) (**b**) Volume of interest quantification of ^89^Zr-Df-BMSC EVs uptake at various timepoints p.i. (n=3). (**c**) *Ex vivo* biodistribution studies of ^89^Zr-Df-BMSC EVs following the final imaging timepoint at 144 h p.i. (n=3) confirmed the prolonged circulation, hepatic clearance, and tumor uptake of the radiolabeled EVs, as also seen in PET.

## Conclusion

PET studies comprehensively studied the biodistribution and pharmacokinetics of EVs derived from various human cell types. We efficiently and stably radiolabeled EVs without altering their physical properties. Our PET studies show labeled EVs had excellent *in vivo* circulation and exhibited parent cell type-dependent biodistribution. Additionally, imaging studies using immunodeficient NSG mice revealed that the biodistribution of EVs was significantly altered compared to that of immunocompetent mice. Proper utilization of PET imaging in the development of EVs should significantly influence and accelerate their clinical translation. Our results demonstrate the significant potential of EVs in diverse therapeutic applications, ranging from cancer therapeutics to radiomitigators, but also the importance of selecting the right cell source and animal model for a given application. Finally, we evidence how noninvasive PET imaging can be a powerful tool in answering these drug development questions, which will hopefully spur additional interest in EV-based technologies.

## Contributions

All authors have contributed to, read, and approved the manuscript. Z.T.R and R.H. conceived the idea and designed the study. Z.T.R., A.S.T., and J.A.K. conducted the experiments. E.A.S. and J.W.E. produced Zr-89. P.H contributed through discussion.

## Ethics Approval

All animal studies were conducted under protocols approved by the Institutional Animal Care and Use Committee at the University of Wisconsin-Madison.

